# Reduced sensorimotor beta dynamics could represent a “slowed movement state” in healthy individuals

**DOI:** 10.1101/2022.02.17.480936

**Authors:** Ryan B. Leriche, Nicholas Jackson, Kathryn Peterson, Zeeya Aspandiar, Vanessa Hufnagel, Nicole C. Swann

## Abstract

Beta oscillations (~13-30 Hz) recorded from the sensorimotor cortex have canonical amplitude changes during movement. Specifically, a movement-related beta decrease (MRBD) occurs before movement, and a post-movement beta rebound (PMBR) follows. We investigated how the MRBD and PMBR vary with movement speed. Individuals performed a task with blocks that generated longer reaction times (RTs) and shorter RTs (Slow and Fast blocks, respectively) while scalp-electroencephalography (EEG) was recorded. The timing of task events before movement was also modulated to generate blocks with certain and uncertain timing (Fixed and Varied blocks, respectively). Beta modulation was reduced in Slow blocks compared to Fast blocks (i.e., a less negative MRBD and less positive PMBR). For the movement certainty manipulation, we saw mixed behavioral and EEG results. Our primary findings align with previous work which has shown reduced movement-related beta modulation in patients with Parkinson’s disease. We propose that a “slowed movement state”, whether it is experimentally induced or a manifestation of Parkinson’s disease bradykinesia, is represented through reduced beta dynamics. Altogether, the MRBD and PMBR may represent motor speed on a continuum with Parkinson’s disease as an extreme example of slowed movement.

## Introduction

Sensorimotor beta oscillations (~13-30 Hz) have been implicated in movement preparation, generation, and termination (see Kilavik et al., 2013 for review). In general, ~500 ms before movement, a movement-related beta decrease (MRBD) occurs, followed by minimal beta power during movement, and for 300-1500 ms after a response, a post-movement beta rebound (PMBR) or beta increase occurs (Alegre et al., 2006; Heideman et al., 2017; Heinrichs-Graham et al., 2016; Muralidharan & Aron, 2021; Tzagarakis et al., 2010). Beta is associated with movement speed in healthy controls (HC)s (Parkes et al., 2006; Zhang et al., 2020) and is pathologically altered in Parkinson’s disease (PD) (Cole et al., 2017; Heinrichs-Graham et al., 2014; Jackson et al., 2019) which is characterized by bradykinesia (i.e. slowed movement). Previous work in HCs has shown a reduced magnitude of the MRBD (i.e., less negative beta) (Heideman et al., 2017; Heinrichs-Graham et al., 2016; Muralidharan & Aron, 2021) and PMBR (i.e., less positive beta) (Parkes et al., 2006; Zhang et al., 2020) during slower compared to faster movements. Similarly, in PD both the MRBD and PMBR have been observed to be reduced in magnitude (Heinrichs-Graham et al., 2014; Meissner et al., 2018; Gert Pfurtscheller et al., 1998; te Woerd et al., 2014, but see Rowland et al., 2015). While beta is related to speed, it is unclear if beta exists on a continuum such that a “slowed movement state” in HCs represents a milder version of the same phenomenon that contributes to parkinsonian bradykinesia. To induce a “slowed movement state” in HCs, we manipulated response demands of a choice GO task. Our hypotheses were that beta power would be higher overall throughout our Slow blocks compared to Fast blocks, but that the relative MRBD and PMBR would be reduced in Slow blocks (Fig. 1).

**Figure 1:**
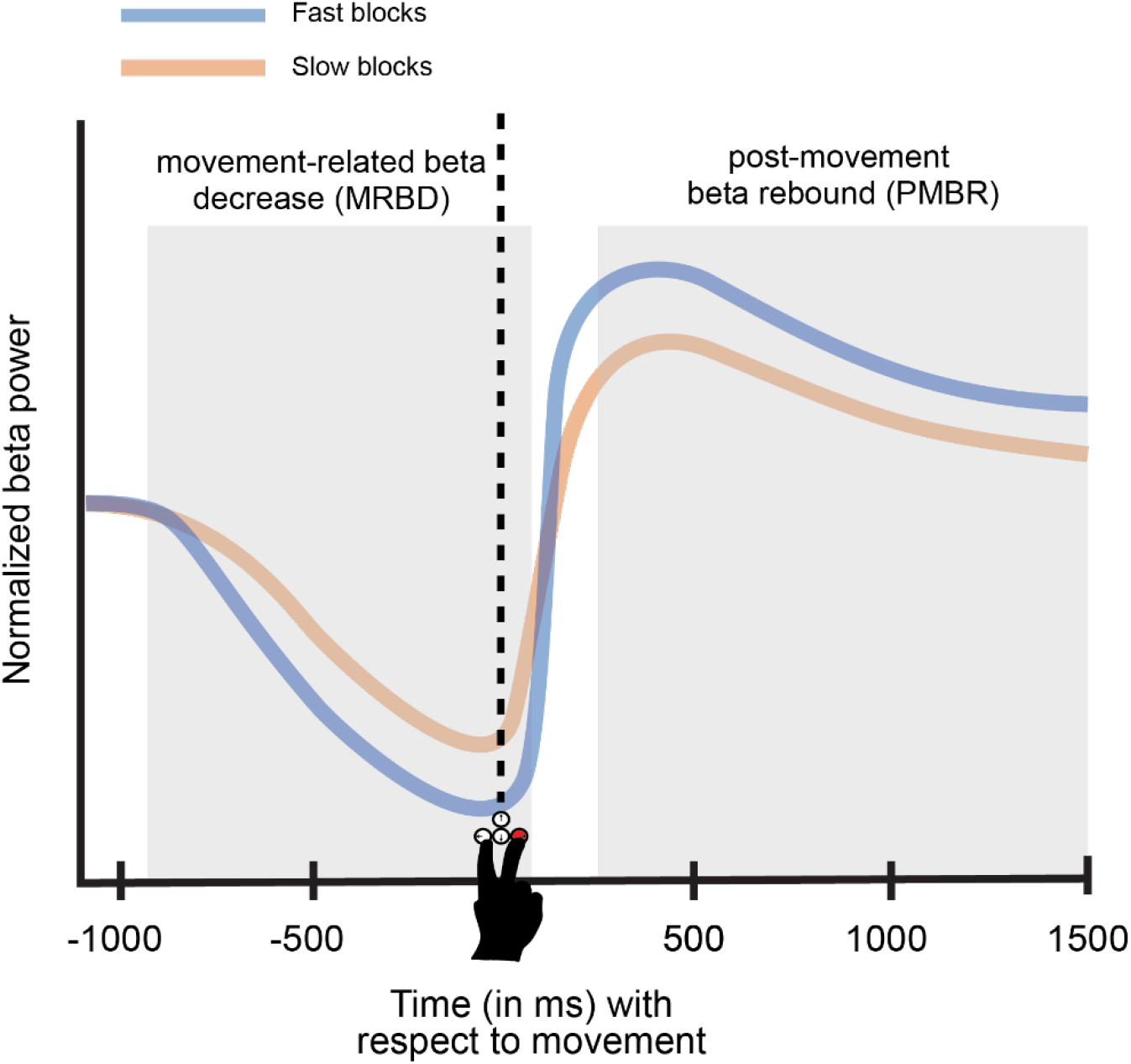
Hypothesized beta activity. Beta power over the contralateral sensorimotor cortex decreases before movement (MRBD) and increases after movement (PMBR) (see Kilavik et al., 2013 for review). This schematic shows beta power over time with movement at 0 ms represented with a vertical dotted line. We expected our Slow blocks to have a MRBD and PMBR of reduced magnitude (compared to Fast blocks) based on what has been observed in PD bradykinetic movement (Heinrichs-Graham et al., 2014; Meissner et al., 2018; Gert Pfurtscheller et al., 1998; te Woerd et al., 2014) and slowed HC movement (Muralidharan & Aron, 2021; Parkes et al., 2006; Zhang et al., 2020). Note that while the normalized (baseline corrected) MRBD and PMBR are shown, we also expected beta to be overall higher throughout Slow blocks.

In addition to speed, we also studied the effects of movement certainty. With greater temporal predictability (i.e., a participant being more certain about *when* they will be instructed to move), a larger MRBD has been observed for foreperiods (FPs) longer than 1 second (Alegre et al., 2006; for directional certainty see Tzagarakis et al., 2010). We investigated if this pattern with regards to temporal certainty is maintained for FPs shorter than 1 second. Our hypotheses with respect to certainty were: (1) FPs with a fixed duration would be more predictable than FPs with a varied duration—and this would be reflected through quicker reaction times in the former and (2) certainty would increase throughout a Varied FP since as time elapses, the probability of GO cue presentation increases. Accordingly, we thought the MRBD magnitude would be increased (i.e., more negative) in Fixed blocks compared to Varied blocks and that within Varied trials, trials with longer FPs would have MRBDs of greater magnitudes since these trials would be associated with greater movement certainty.

## Methods

### Participants

All participants provided written informed consent in accordance with the Institutional Review Board of the University of Oregon and the Declaration of Helsinki. Subjects were recruited for the study via flyers around the University of Oregon’s campus and via the University of Oregon’s online experiment participation portal (SONA). These participants were paid $10-12/hour for their participation. The eligibility for the study included: 18-40 years old, not diagnosed with a movement or motor impairment/disorder, no neurological disorders, not taking neurological or psychiatric medications (including those for depression or ADHD), fluent in English, right-hand dominance, and normal or corrected-to-normal vision. The study was comprised of one ~1.5-hour EEG recording session. We tested 14 participants (11 females, 3 males). Two participants were later excluded from the analysis due to EEG data quality (detailed in the EEG preprocessing section), so a total of 12 participants’ data were included in the final analysis. All values are reported as mean ± standard deviation. Participants were on average 21.33 ± 3.35 years old (range 18 to 29 years old).

### Task design

A blocked-design choice GO task was implemented while participants sat approximately two feet away from a 21-inch computer monitor within a Faraday cage. They used their right index finger and right middle finger to press the left and right arrows keys of a response keypad, respectively. Each trial consisted of a set cue presented during a foreperiod (FP), a GO cue during a response window (RW), and finally an inter-trial interval (ITI) (Fig. 2). The set cue was a white square in the center of the screen while the GO cue was a white right or left facing arrow—presented with equiprobability—indicating which arrow key to press. Subjects were instructed to respond during the RW (i.e., before the GO cue disappeared). Two variables were manipulated between blocks: the duration of the FP (Fixed and Varied) and the duration of the RW (Slow and Fast), leading to 4 block-types (Slow/Fixed, Fast/Fixed, Slow/Varied, and Fast/Varied, Fig. 2C). In Fixed blocks, the FP was always 500 ms, while in Varied blocks the FP was randomly selected to be 300, 400, 500, 600, or 700 ms per trial. Each participant’s mean reaction time (RT) was calculated based on a practice session of 20 trials completed just before the experiment commenced with a Varied FP and RW of 800 ms (Fig. 2A). For Fast blocks, the RW was set as the mean RT. In contrast, the RW for the Slow blocks was four times the mean RT (Fig. 2B). For all trials after the RW, a blank screen was presented during the ITI which lasted 2300 to 2700 ms. This behavioral task was created with MATLAB (release R2020a) using the Psychophysics Toolbox Version 3 functions (Brainard, 1997; Kleiner et al., 2007; Pelli, 1997).

**Figure 2.**
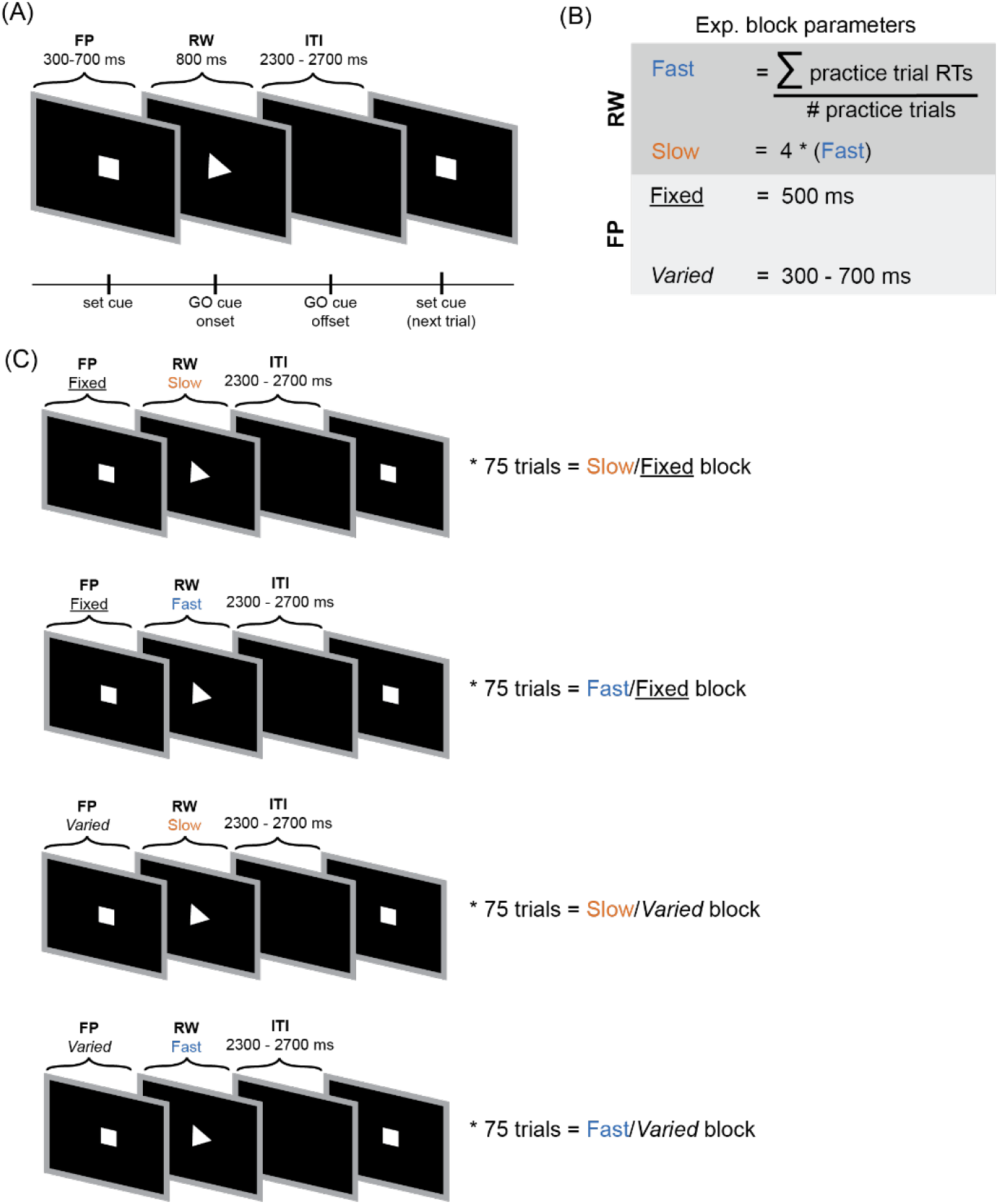
Movement speed and certainty task schematic. (A) Participants first completed 20 practice trials with a 300 - 700 ms foreperiod (FP), an 800 ms response window (RW) and 2300 - 2700 ms inter-trial interval (ITI). (B) From the practice trials, the timing parameters for the experimental blocks were calculated. (C) Participants responded to four block types (Slow/Fixed, Fast/Fixed, Slow/Varied, and Fast/Varied) all of which were 75 trials long shown in four respective timelines. Each block was presented twice in a randomized order totaling 600 trials over 8 blocks.

Participants completed 8 blocks of 75 trials (600 trials total) with each of the four experimental blocks presented twice in a random order. At the end of every block, participants could take a break and indicated when they were ready to continue. During this break, the percentage of correct trials in the last block was shown to incentivize task performance. Trials were considered incorrect if participants: (1) responded within 100 ms of the GO cue presentation—to minimize premature responses, (2) after the RW, and (3) if the wrong keyboard arrow was pressed.

### EEG recording and preprocessing

Sixty-four channel EEG was recorded with a BioSemi ActiveTwo system sampled at 1024 Hz. Additional electrodes were placed on the left and right mastoid, lateral to each eye, below the right eye (to monitor for eye movements and blinks), and on the right extensor carpi radialis longus to record electromyography. Note that electromyography was not analyzed here.

EEG data was preprocessed using custom MATLAB scripts and the EEGLAB toolbox (Delorme & Makeig, 2004) in-line with previous work (Swann et al., 2015). Each individual channel was re-referenced to the common average and slow signal drifts were removed with a 0.5 Hz high pass filter. Data were then decomposed using an Independent Component Analysis, and components associated with saccades and blinks were removed from the data through manual inspection using the ‘pop_runica’ function in EEGLAB (Delorme et al., 2007). The remaining components were backpropagated into channel space. Two of the 14 participants’ data were excluded because of unidentifiable blink and/or saccade components—meaning 12 subjects’ data were used for subsequent EEG and behavioral analyses. The EEGLAB functions ‘pop_jointprob’ and ‘pop_rejkurt’ were then used to reject epochs with absolute values, and/or kurtosis values, that were 5 standard deviations above their respective means. On average, 79.18 ± 7.61% of trials per subject were included for subsequent analyses.

### Spectral analysis

Due to our interest in motor activity, our primary analysis focused on channel C3 which corresponds most closely to left sensorimotor cortex (contralateral to the responding right hand). With the EEGLAB function ‘spectopo’, we generated a power spectrum in decibels (dB =10* log_10_(μV^2^ /Hz)) per each experimental block by finding the power spectrum per trial and averaging across trials within condition per each subject. To evaluate event-related beta modulation, we generated event-related spectral perturbations (ERSP)s from C3 using a two-way FIR1 filter (EEGLAB function ‘eegfilt’) with a 3 Hz bandwidth from 4 - 64 Hz. A Hilbert transform was then applied to the filtered signal to generate a complex signal. Squaring the modulus of this complex signal gave us the power of our signal. We then used a single-trial z-score normalization which has been shown to be more noise resistant than classical additive and gain models of EEG baselines (Grandchamp & Delorme, 2011). In brief, over the entirety of a single trial, that trial’s mean and standard deviation of power was calculated per frequency. Each time-frequency point was then subtracted from the mean value for that particular frequency and divided by the standard deviation for the specific frequency. This produced a time-frequency z-score matrix per trial. Per experimental block, each subject’s data were then averaged across trials. Next, the z-scored ERSPs per condition were averaged across subjects. The final output represents the average per condition over all subjects and trials—a grand averaged ERSP. To visualize beta changes more easily, we also created so-called beta traces by averaging over the beta band (13-30 Hz) to generate a vector of mean-beta values over time.

While our *a priori* analysis plan focused on C3, as an exploratory analysis we examined beta power over all channels. Guided by movement-related beta changes from our initial C3 analysis, we examined beta from −250 to 250 ms (with movement at 0 ms) to examine the MRBD and 250 to 750 ms to examine the PMBR. Notably, 250 to 750 ms was selected to examine the PMBR to minimize visual confounds between blocks. The GO cue disappeared at ~1200 ms on average after movement in Slow blocks and disappeared much earlier in Fast blocks (with exact values for each dependent on an individual subject’s mean RT). Thus, the 250 to 750 ms timeframe was selected since it didn’t include a *change in visual stimuli* for either block. During the PMBR in Fast blocks, the GO cue would have already disappeared and in Slow blocks the GO cue would be continuously presented. Additionally, we examined 250 ms to 750 ms before movement to examine preparatory changes during the FP. Finally, we included 1000 to 500 ms before the set-cue (i.e., during the ITI) as a control comparison. By averaging across the beta band and our timeframes of interest, we explored beta changes over the entire scalp topography throughout our task epochs.

### Statistical Analyses

Repeated measures ANOVAs were performed to compare across multiple levels for our one-dimensional data. To test specific hypotheses, Tukey’s multiple comparison test (implemented with MATLAB’s ‘multcompare’ function) was used.

For group-level analysis to assess ERSPs, we used a nonparametric cluster-based permutation test—which corrects for multiple comparisons by clustering over adjacent time and frequency points—to compare between conditions within the same subject (Maris & Oostenveld, 2007). The same approach was used for our beta traces based on temporal adjacency. Likewise, for our scalp-topographies (the mean beta power across channels per timeframe), clustering was based on channels being adjacent in physical space with a minimum cluster size of two channels. Since our topography analysis was exploratory, we used a Bonferroni correction to determine significant clusters given our four timeframes of interest (i.e., permutation p-values < 0.0125 (0.05 ÷ 4) for a cluster to be considered significant). For all cluster-based approaches, we used the FieldTrip toolbox function ‘ft_freqstatistics’ (Oostenveld et al., 2011).

## Results

### RTs were altered as a function of movement speed and possibly by movement certainty at the block level

Overall, participants performed well with a mean RT of 341 ± 26 ms (range 296 to 375 ms) and mean accuracy of 93.5 ± 4.0% (range 84.5 to 97.7%). A repeated measures 2-way ANOVA was used to evaluate the effects of movement speed and certainty across the subjects’ mean RTs and response accuracy. We saw a main effect of movement speed on RT (p = 1.39 * 10^−6^). Tukey’s multiple comparison test then confirmed that Slow blocks had longer group-mean RTs (370 ± 34 ms) than Fast blocks (325 ± 22 ms) (p = 1.39 * 10^−6^). There was no effect of movement certainty on RT (p = 0.931) or an interaction effect (p = 0.17). Fixed and Varied blocks had a group-mean RT of 349 ± 29 ms, and 349 ± 27 ms, respectively. For response accuracy, we found a significant effect of movement speed (p = 0.0002), but not of movement certainty (p = 0.500) or an interaction effect (p = 0.800). A follow-up Tukey’s test showed participants responded more accurately in Slow blocks (99.45 ± 0.45%) than Fast blocks (87.49 ± 7.84%) (p = 0.0002). Fixed and Varied blocks had a mean response accuracy of 93.94 ± 5.21%, and 93.03 ± 3.95%, respectively.

The decreased accuracy of Fast blocks compared to Slow blocks was likely due to the more stringent RW length in Fast blocks leading to many responses after the RW. For Fast blocks, 11.25 ± 7.63% (range = 3.67 to 28.28%) of trials did not have responses during the RW while only 0.33 ± 0.35% (range = 0 to 1.65%) of Slow block responses did not occurring during the RW. Since trials with late or no responses were considered *incorrect* and these RTs were not recorded, this created a confound in which the slower end of our Fast block RT distribution was artificially cut off. To correct for this, for each subject we took whatever percentage of Fast trials were missed and removed that same percentage of the “slowest” Slow trials per subject (e.g., if 10% of Fast trials were missed, only the fastest 90% of Slow trials were kept).

With the “slowest” Slow trials removed, our behavioral results were similar. Our main effect of movement speed on RT was maintained (*p* = 2.14 * 10^-5^). Tukey’s test showed that our corrected Slow blocks still had longer group-mean RTs (356 ± 31 ms) than Fast blocks (325 ± 22 ms) (*p* = 2.14 *10^-5^) (Fig. 3B). We still saw no effect of movement certainty on RT (*p* = 0.627). The corrected Fixed and Varied blocks had a group-mean RT of 340 ± 27 ms, and 341 ± 25 ms, respectively. However, we found a trending interaction effect between movement speed and certainty (*p* = 0.0659). As an exploratory investigation we ran a paired t-test, finding that Fast/Varied blocks had longer a group-mean RT than Fast/Fixed blocks (329 ± 23 ms, and 321 ± 22 ms, respectively, *p* = 0.0142)). Overall, our approach of counting late RTs as incorrect did not drive our main observed behavioral effect. For consistency, the Slow trials removed from this corrected behavioral analysis were also removed from the EEG analysis.

**Figure 3.**
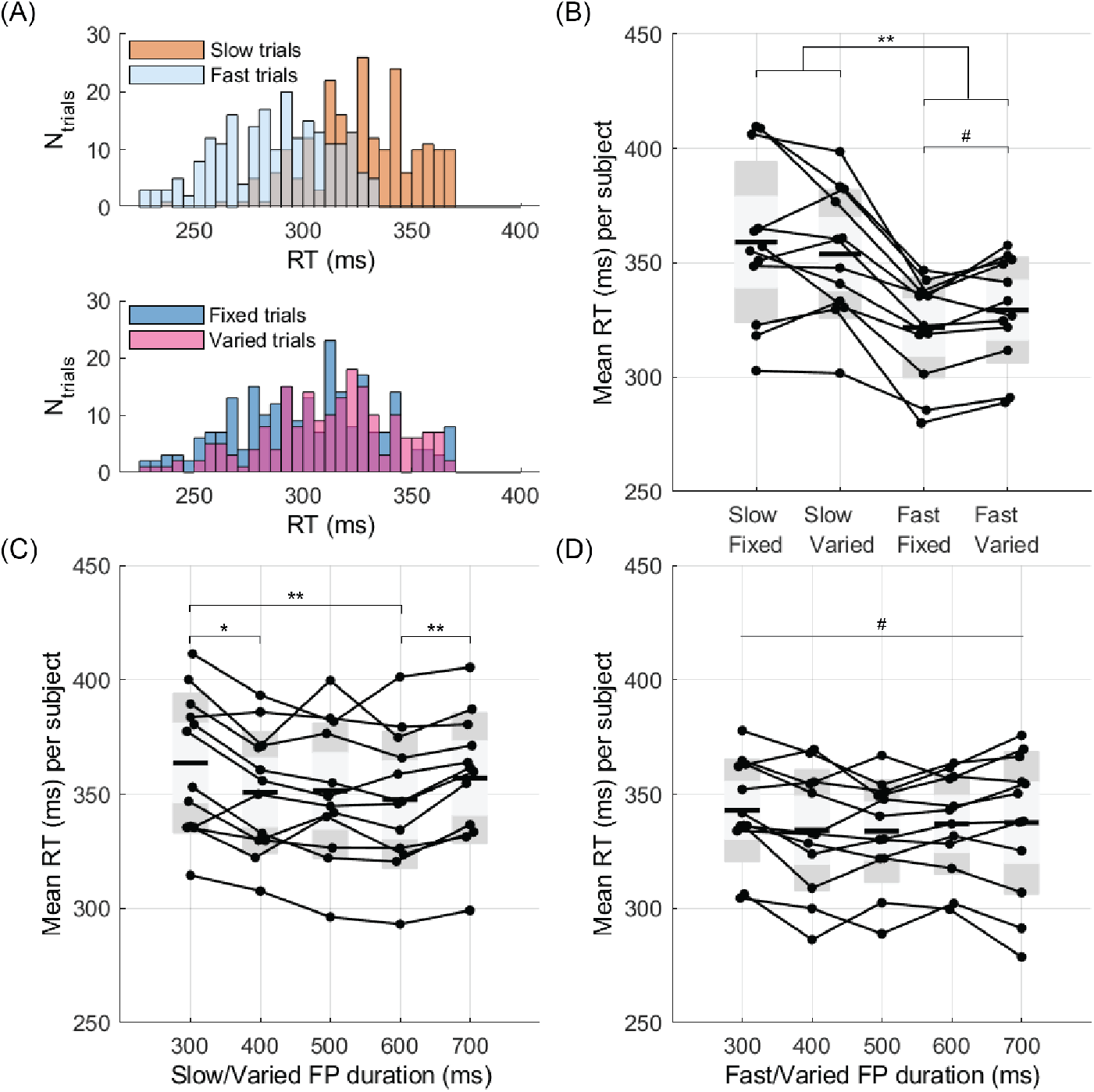
RTs were manipulated by movement speed blocks and marginally by certainty blocks. (A) An example subject’s RTs. RTs were qualitatively longer in Slow compared to Fast trials, but no meaningful difference was seen between Fixed and Varied trials. (B-D) Each point is single subject’s mean RT over the specified block type with lines used to connect mean RTs from the same subject. (B) Participants had longer RTs for Slow blocks than Fast blocks (*p* = 2.14 * 10^-5^), but no difference between Fixed and Varied blocks (*p* = 0.627). However, we found a trending interaction effect with Fast/Fixed blocks having shorter RTs than Fast/Varied blocks (*p* = 0.0659). (C) Across the FPs (300 - 700 ms) of Slow/Varied trials, 300 ms FP trials had longer group-mean RTs than 400 ms and 600 ms FP trials (*p* = 0.02 and *p* = 0.001, respectively). The 700 ms FP trial also had a longer group-mean RT than the 600 ms FP trial (*p* = 0.001). (D) For the Fast/Varied trials, there was a trending difference in RT across FP trials (*p* = 0.0506) with 300 ms FP trials having the longest RT. #p< 0.1; *p< 0.05; **p < 0.001.

While we only saw trending RT effects between movement certainty blocks, we also examined if the mean RT changed with FP duration within Varied blocks— hypothesizing that movement certainty would increase as the FP went on, reflected through quicker RTs in longer FP trials (Fig. 3C & 3D). A repeated measures one-way ANOVA showed a trending difference between the subjects’ mean RTs across the 300, 400, 500, 600, and 700 ms FPs of Fast/Varied trials (*p* = 0.0506) with group-mean RTs of 343 ±23 ms, 334 ± 27 ms, 334 ± 23 ms, 337 ± 22 ms, and 337 ± 31 ms, respectively. Notably, the 300 ms Fast/Varied FP trials had the longest group-mean RT. A repeated measures one-way ANOVA of the different FPs of Slow/Varied trials revealed a significant effect of FP on RT (*p* = 0.0002). A follow-up Tukey’s test revealed that the Slow/Varied FP trials with a 300 ms FP had a longer group-mean RT than both the 400 ms FP trials (*p* = 0.02) and 600 ms FP trials (*p* = 0.001) (364 ± 31 ms versus 351 ± 27 ms and 348 ± 30 ms, respectively). The 700 ms FP Slow/Varied trials also had a longer group-mean RT (357 ± 29 ms) than the 600 ms Slow/Varied FP trials (348 ± 30 ms) (*p* = 0.001). The 500 ms Slow/Varied FP trials had a group-mean RT of 351 ± 30 ms. Altogether, this provides weak statistical evidence that the shorter Varied FPs were less certain than the longer Varied FPs.

### Grand averaged ERSPs and beta traces reveal reduced alpha/beta dynamics in Slow blocks

To compare movement speed blocks, we first examined the ERSPs over C3 (Fig. 4). As expected, there was a prominent beta (along with an alpha (9-12 Hz)) desynchronization at the time of movement followed by a rebound, as is typically observed over sensorimotor channels (Kilavik et al., 2013; G. Pfurtscheller, 1981). Slow/Fixed blocks had increased 9-17 Hz power from −405 to 240ms, with respect to movement, compared to Fast/Fixed blocks (*p* = 0.004) and decreased 4-16 Hz power from 253 to 1699 ms after movement (*p* = 5 * 10^-4^). This shows a greater beta/alpha desynchronization before movement and a greater rebound in Fast/Fixed blocks compared to Slow/Fixed blocks. Similarly, after movement, Slow/Varied blocks had elevated 9-20 Hz power from 60 to 462 ms (*p* = 0.009) and decreased 9-25 Hz power from 484 to 1515 ms (*p* = 0.0030). Finally, Slow/Varied blocks also had elevated 4-8 Hz power from 1455 to 2150 ms (*p* = 0.0180) and elevated 9-25 Hz power from 1844 to 2150 ms (*p* = 0.0195) compared to Fast/Varied blocks. The ERSPs of C3 demonstrate that both Slow blocks were associated with a lesser PMBR and that the Slow/Fixed block also showed a reduced MRBD.

**Figure 4.**
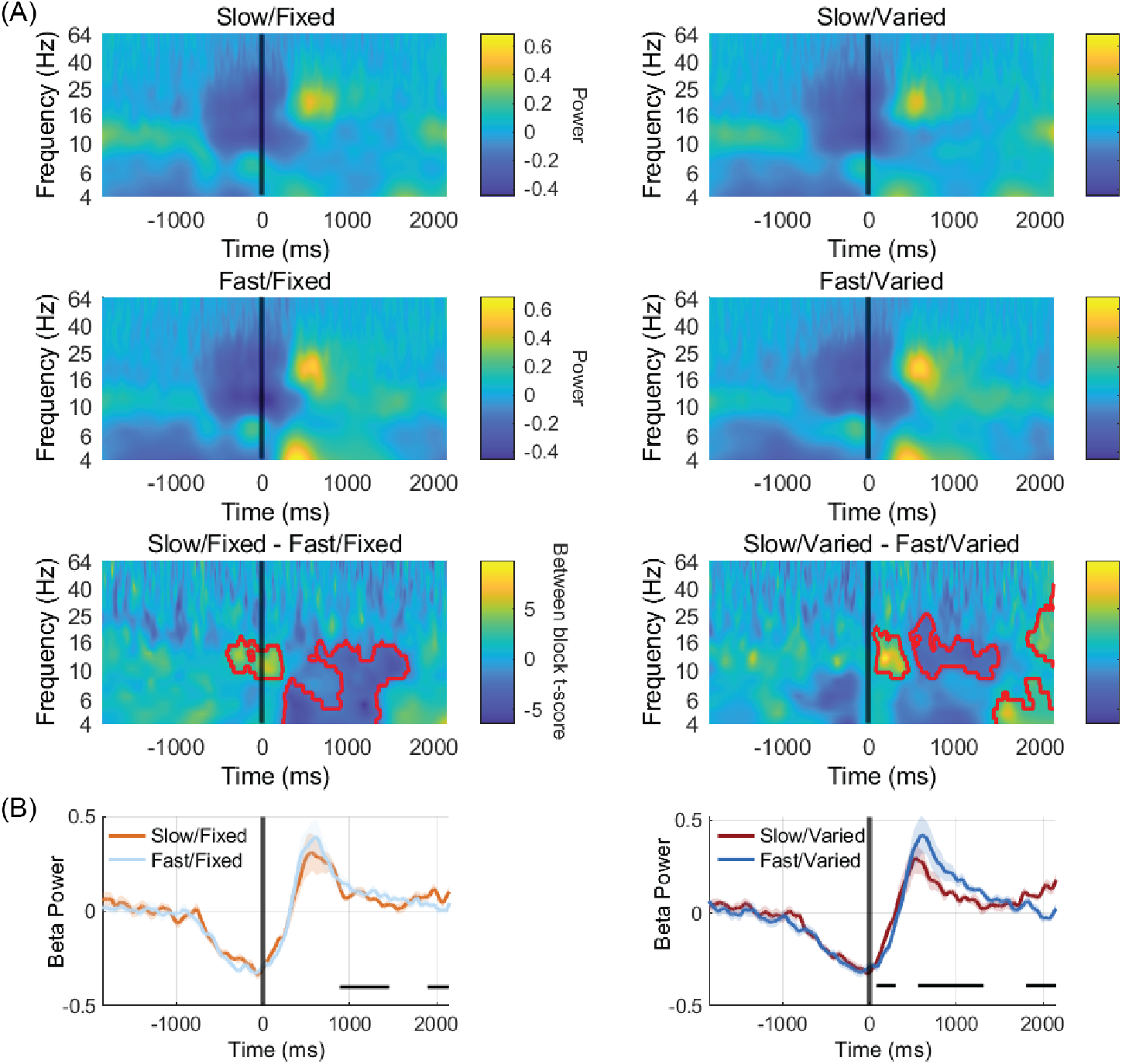
Grand averaged ERSPs and beta traces over sensorimotor cortex showed that Slow blocks had a diminished post-movement beta rebound (PMBR) compared to Fast blocks. (A-B) All plots are from channel C3. (A) The grand averaged ERSPs for Slow/Fixed, Fast/Fixed, Slow/Varied, and Fast/Varied blocks are in the first two rows. Each exhibits canonical beta activity around movement, but with different magnitudes. In the bottom row, significant time-frequency clusters between experimental blocks are outlined (see text for discussion). (B) The grand averaged beta traces (13-30 Hz mean over time) per experimental block. Significantly different time periods are indicated with horizontal black bars below the traces and shading representing the SEM.

As our *a priori* hypotheses focused on beta, we also averaged across the beta range (13-30 Hz) to clearly show the changes relative to movement for each condition (Fig. 4B). The Slow/Fixed beta trace had a higher amplitude than the Fast/Fixed beta trace from 1900 to 2145 ms (*p* = 0.012), but a lower amplitude from 888 to 1450 ms (*p* = 0.0015). Similarly, Slow/Varied blocks had elevated beta power compared to Fast/Varied blocks from 82 to 304 ms (*p* = 0.020) and from 1810 to 2150 ms (*p* = 0.0025) but decreased beta power from 560 to 1319 ms (*p* = 0.001) (Fig. 4B). Since our movement certainty blocks did not show consistent behavioral effects, we focused our EEG analysis on the Slow and Fast blocks. All the results discussed here were corrected with a cluster-based approach to account for multiple comparisons (see Statistical Analyses under Methods). Regardless of movement certainty, the beta traces showed that the PMBR was reduced in Slow blocks compared to Fast blocks. Overall, our ERSPs and beta traces demonstrated a diminished sensorimotor PMBR in Slow blocks. There was also a reduced MRBD which was mainly evident in the ERSPs, and only briefly apparent for the beta traces (in the Slow/Varied vs Fast/Varied comparison). The discrepancy between the ERSPs and the beta traces was likely due to both averaging across the entire beta band in the beta trace plots and also the inclusion of lower frequencies (below 12 Hz) in the ERSP plots. For simplicity, we’ll continue to refer to these effects as the MRBD and PMBR but note the differences between Slow/Fixed and Fast/Fixed ERSPs were alpha dominate and spanned down as low as 4 Hz at times.

### The power spectrum from C3 was similar between experimental blocks

To see if the observed differences in beta were due to a shift in the oscillatory activity throughout the entirety of a block, we examined the power spectrum from C3 per subject (Fig. 5). While there was inter-subject variability, no systematic differences were observed between experimental blocks as shown in the grand averaged power spectrum (Fig. 5A). To derive a mean beta value per subject, we averaged over the beta range to obtain one beta value per subject for each block (Fig. 5B). A repeated measures two-way ANOVA showed no difference in the mean beta power between Slow (−5.40 ± 2.16 dB) and Fast blocks (−5.46 ± 2.42 dB) (*p* = 0.683), nor Fixed (−5.35 ± 2.43 dB) and Varied blocks (−5.52 ± 2.19 dB) (*p* = 0.4090), but there was a trending interaction effect (*p* = 0.0786). For this trending interaction, Slow/Varied blocks had higher beta power (−5.30 ± 2.06 dB) than Fast/Varied blocks (−5.87 ± 2.47 dB). This suggests that the *reduced* PMBR of Slow/Varied blocks seen in our ERSPs may have been partly due to elevated beta throughout the entirety of Slow/Varied blocks.

**Figure 5.**
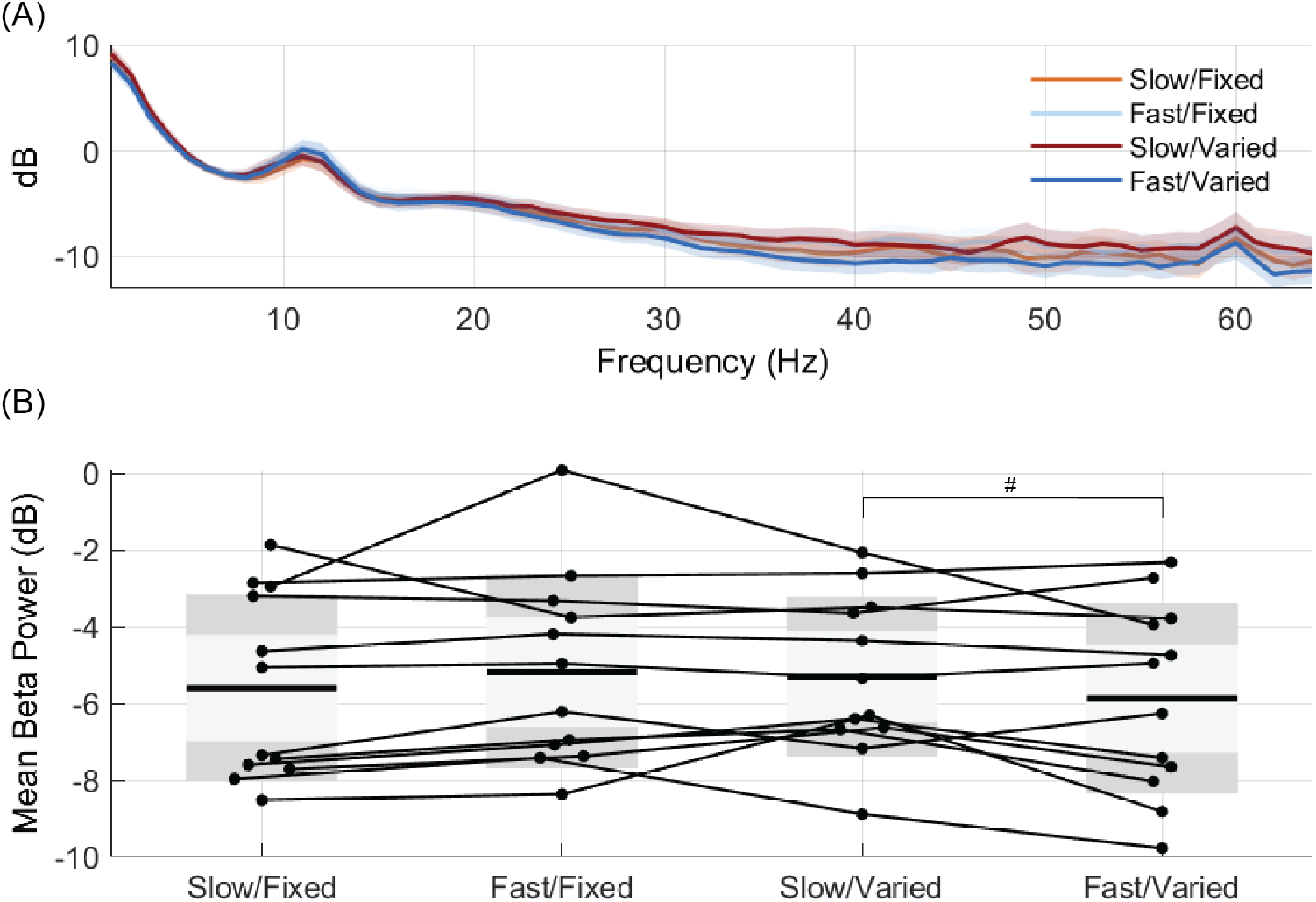
Minimal power spectra differences between blocks over C3. (A) The grand-averaged power spectrum of all subjects per experimental condition (with the SEM shown as shading). (B) The mean beta power across all time points per subject showed no main effects, but a trending interaction of Slow/Varied blocks having elevated beta power than Fast/Varied blocks (*p* = 0.0786). The lines connecting experimental conditions represent data from the same subject. #p< 0.1.

### An exploratory topography analysis revealed reduced beta modulation in Slow blocks over frontal, central, and parietal areas

After observing reduced beta dynamics over C3, we explored beta across all channels. As described above, we had four epochs of interest: (1) 1000 to 500 ms before the set cue, (2) −750 to −250 ms, relative to movement at time 0, (3) −250 to 250 ms, again movement at time 0, and (4) 250 to 750 ms with movement at time 0. By averaging across the beta band and within the four epochs, we saw reduced beta modulation in Slow blocks over frontal, central, and parietal channels (Fig. 6; Table 1 for exact channels). During movement, Slow/Fixed blocks had a reduced MRBD (i.e., less negative beta) than Fast/Fixed blocks over frontal, central, and parietal channels (*p* = 5 * 10^-4^). Similarly, Slow/Varied blocks had a reduced MRBD than Fast/Varied blocks over frontal, central, and parietal channels (*p* = 5 * 10^-4^). After movement, Slow/Fixed blocks had a reduced PMBR (i.e., less positive beta) than Fast/Fixed blocks over frontal, parietal channels, and central channels (*p* = 0.0015). A similar effect was seen for Slow/Varied blocks versus Fast/Varied blocks (*p* = 0.0080). During the FP, Slow/Varied blocks had more positive beta power compared to Fast/Varied blocks—suggesting that the Varied FP manipulation interacted with Slow blocks; however, this was not apparent in the behavioral data. Notably, C3 was not significantly different for the PMBR comparisons nor the MRBD of Slow/Fixed and Fast/Fixed comparison—likely due to data being averaged across the entire timeframe. Despite this, our topography analysis maintained the reduced PMBR effect seen in the ERSPs of C3, but also provides additional support that participants were in a “slowed movement state” since the MRBD was diminished over more widespread cortical areas.

**Figure 6.**
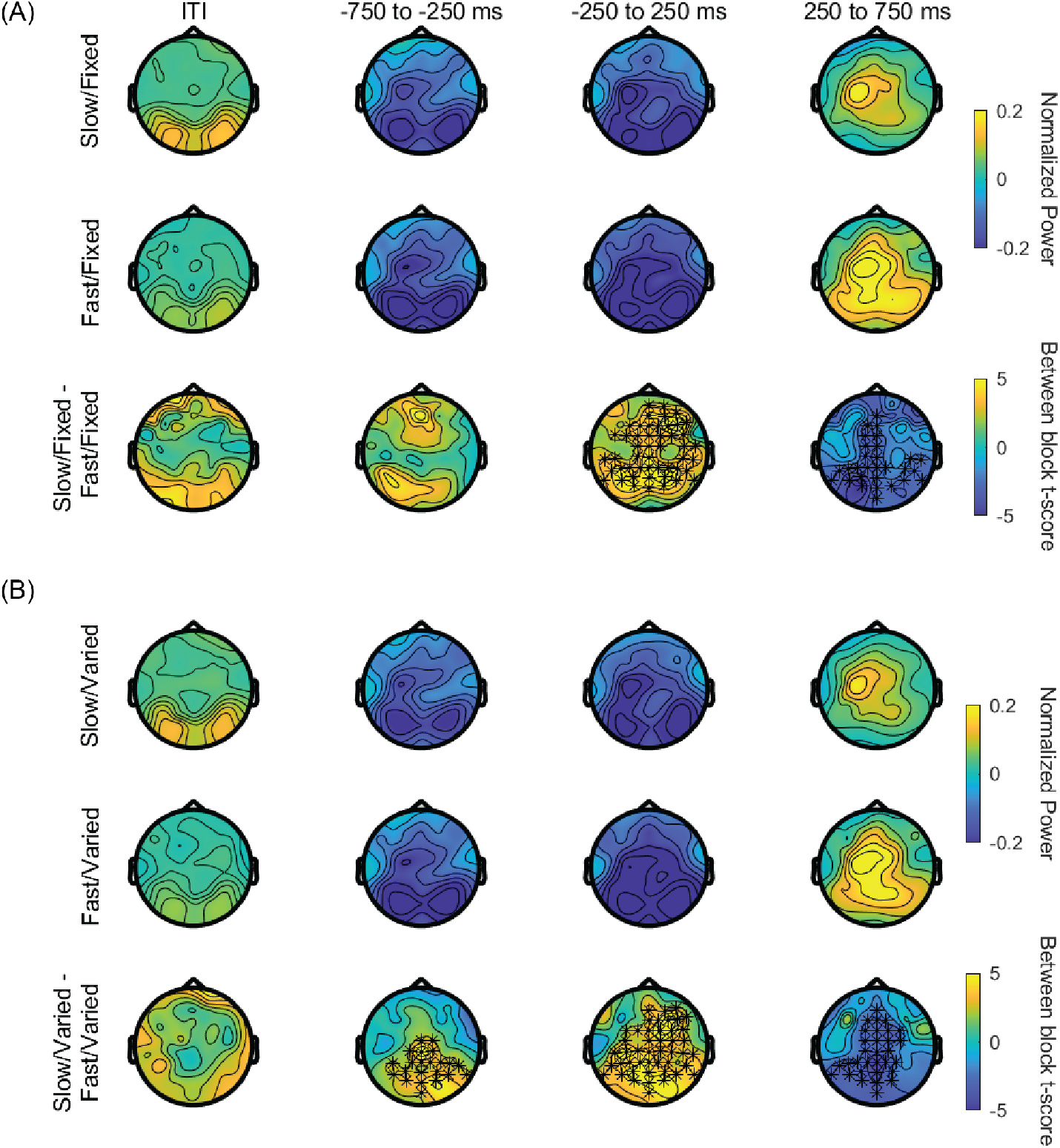
Over frontal, central, and parietal channels Slow blocks had reduced beta modulation. (A-B) The time frames from the left to right columns are: −1000 to −500 ms (set cue at 0 ms, during the ITI), and with movement at 0 ms: −750 to −250 ms, −250 to 250 ms, and 250 to 750 ms. These time frames were motivated from the MRBD and PMBR observed in the C3 analysis. The top two rows of sub-panels (A) and (B) show the grand averaged beta band topographies during the Slow and Fast blocks per Fixed blocks (A) and Varied blocks (B). The bottom row per sub-panel shows the difference in mean beta between Slow and Fast blocks, within a timeframe, with significant channels starred. *p < 0.0125.

**Table 1.** Significant channels per timeframe from topography analysis (multiple comparison corrected)

| Comparison | ITI | FP | Movement (MRBD) | Post-movement (PMBR) |
| --- | --- | --- | --- | --- |
| Slow/Fixed versus Fast/Fixed | None | None | F1, F3, FC5, FC3, FC1, C1, TP7, CP5, CP1, P1, P3, P5, P7, P9, PO7, PO3, O1, Oz, POz, Pz, CPz, Fpz, Fp2, AF8, AF4, AFz, Fz, F2, F4, F6, F8, FC6, FC4, FC2, FCz, Cz, C2, C6, T8, TP8, CP6, P2, P4, P6, P8, P10, PO8, PO4, O2<br>($p = 0.0005$) | F1, FC1, C1, CP1, P1, P3, P5, P7, P9, PO7, PO3, O1, Iz, Oz, POz, Pz, CPz, AFz, Fz, FCz, Cz, TP8, CP6, P2, P6, P8, P10, PO8, O2<br>($p = 0.0015$) |
| Slow/Varied versus Fast/Varied | None | CP1, P1, P5, PO7, PO3, O1, Iz, Oz, POz, Pz, CPz, Cz, C2, CP2, P2, P4, P6, P8, P10, PO8, PO4, O2<br>($p = 0.0025$) | FC3, FC1, C3, CP5, CP3, CP1, P1, P3, P5, P7, P9, PO7, PO3, O1, Iz, Oz, POz, Pz, CPz, AF8, AF4, AFz, Fz, F2, F4, F6, F8, FC6, FC4, FC2, FCz, Cz, C2, C4, C6, TP8, CP6, CP4, CP2, P2, P4, P8, P10, PO8, PO4, O2<br>($p = 0.0005$) | F1, FC1, C1, CP1, P1, P3, P5, P7, P9, PO7, PO3, O1, Iz, Oz, POz, Pz, CPz, AFz, Fz, F2, FC4, FC2, FCz, Cz, C2, C4, CP2, P2, P4, PO4, O2<br>($p = 0.0080$) |
| Sig. in Fixed, but not Varied comparison | N/A | N/A | C1, F1, F3, FC5, Fp2, Fpz, P6, T8, TP7 | CP6, P10, P6, P8, PO8, TP8 |
| Sig. in Varied, but not Fixed comparison | N/A | CP1, P1, P5, PO7, PO3, O1, Iz, Oz, POz, Pz, CPz, Cz, C2, CP2, P2, P4, P6, P8, P10, PO8, PO4, O2 | C3, C4, CP2, CP3, CP4, Iz | C2, C4, CP2, F2, FC2, FC4, P4, PO4 |

The leftmost column shows which conditions the significant channels are from. The next four columns from left to right show data from four timeframes: −1000 to −500 ms (set cue at 0 ms, during the ITI), and −750 to −250 ms, −250 to 250 ms, and 250 to 750 ms (the latter three with movement at 0 ms). Each cell within the table shows the significant channels for its respective timeframe and condition.

## Discussion

Here, we report that the PMBR was diminished in our Slow blocks compared to Fast blocks—suggesting the PMBR may scale with movement speed on a continuum. While the MRBD was only reduced in the ERSPs of Fixed/Slow blocks, our scalp-topographies showed both a diminished PMBR and MRBD which was widespread and included general sensorimotor regions during our Slow blocks. Our movement certainty manipulation of the FP did not show consistent significant behavioral effects.

This study builds upon the efforts to directly probe movement speed and alter beta activity. A reduced MRBD with longer RTs has been seen by embedding a secondary movement within the previous PMBR (Muralidharan & Aron, 2021), and by inducing a high beta state in macaque monkeys via neurofeedback (Khanna & Carmena, 2017). Likewise in HCs, suppressing beta burst activity with neurofeedback is associated with shorter RTs (He et al., 2020). Our study complements these efforts as we attempted to put participants into a “high beta state” by encouraging slower movements (at the block level) by reducing response urgency. In a similar fashion, Zhang and colleagues explicitly instructed slower and faster movements at the trial level and observed a reduced PMBR during their slower trials.

One puzzle is how a reduced PMBR after slower trials fits with previous findings of the excessive beta synchrony seen in the cortico-basal ganglia loops of PD (Brittain et al., 2014). The distinction may be that high beta synchrony (i.e., elevated beta waveform shape metrics and phase-amplitude coupling) and high beta power are not necessarily concordant. While high beta synchrony has been shown in the cortex of patients with PD, resting sensorimotor beta power shows inconsistent effects (Cole et al., 2017; De Hemptinne et al., 2013; Geraedts et al., 2018; Jackson et al., 2019; Swann et al., 2015). Thus, it may be that the high beta synchrony associated with slowed movement constrains physiological fluctuations of beta power. This possibility is supported by the findings that patients with PD have shown a reduced sensorimotor beta activity throughout task-related movements (Heinrichs-Graham et al., 2014; Meissner et al., 2018; te Woerd et al., 2014, but see but see Rowland et al., 2015), self-initiated movements (Gert Pfurtscheller et al., 1998), and proprioceptive stimulation (Vinding et al., 2019). Here we show that this may extend to HCs under less urgent task demands. While we predicted higher beta power throughout Slow blocks—thus reducing the relative PMBR and MRBD—we only found trending effects. Again, this may be because beta power is not sensitive to all types of beta synchrony associated with slowed movement.

Another explanation for our findings is that the PMBR scales with the information contained within the previous motor plan. In the pre-frontal cortex, since a beta increase is thought to “clear out” information in working memory tasks, the sensorimotor PMBR may have a similar function in motor tasks (Schmidt et al., 2019). Given our Fast blocks had a shortened RW, participants likely were in an enhanced attentional state— corresponding to a more urgent motor plan and thus an enhanced PMBR. However, with a lengthened RW in Slow blocks, participants could rely on a less urgent motor plan and therefore a reduced PMBR. Considering bradykinesia and rigidity in PD, perhaps their high beta synchrony diminishes the ability to form comprehensive motor plans which slows movement and reduces both MRBD and PMBR.

We have demonstrated that a reduced PMBR (and to some extend a reduced MRBD) is associated with the “slowed movement state” induced in our task. Our results in conjunction with previous work in PD (Heinrichs-Graham et al., 2014; Meissner et al., 2019; Gert Pfurtscheller et al., 1998; te Woerd et al., 2014), suggest that slowed movement may exist on a continuum with PD bradykinesia—reflected in the magnitude of the PMBR. Future studies could directly compare beta modulation between the slowed movement of HC to PD patient movement. Further, paradigms to intentionally slow movement should assess how phase-amplitude coupling and non-sinusoidal oscillations measure “slowed movement states” in HCs to determine the specificity of these PD biomarkers (Khanna & Carmena, 2017; Muralidharan & Aron, 2021; Zhang et al., 2020).

### Limitations

Our attempt to alter movement certainty had unclear behavioral effects. We only saw a trending relationship of Fast/Fixed blocks having shorter RTs than Fast/Varied blocks. Perhaps if Fixed and Varied blocks differed more (for instance with Fixed FP of 600 ms and Varied FP of 200-1000 ms rather than Fixed FP of 500 ms and Varied FP of 300-700 ms), a definitive effect may have emerged. Notably, this trending effect coincided with Fast blocks—which had more urgent response demands—possibly suggesting that the temporal predictability of the FP was utilized under more taxing task conditions. This hypothesis is supported by the findings of temporal expectation aiding the utilization of other task parameters to improve performance (Nobre & Van Ede, 2017). The Slow/Varied and Fast/Varied trials with 300 ms FPs had delayed RTs compared to their respective longer FP trials—providing weak evidence that the FP became more certain overtime. While we expected RT to decrease as the FP elapsed in Varied trials, early work suggests that such effects may only be evident for particular FP duration ranges (Niemi & Risto, 1981). Overall, with greater stratification of FP durations, behavioral results may have emerged.

Our ERSPs from channel C3 and our scalp topographies did not display entirely consistent effects. While C3 ERSPs showed a reduced PMBR and MRBD in Slow blocks, the topographies did not reveal a difference in beta modulation for C3. We speculate that this difference may be because the topographies utilized averaged data over the whole beta range and time intervals.

### Conclusion

We observed reduced movement-related beta modulation for Slow compared to Fast blocks. This observation aligns with previous work demonstrating a reduced beta dynamics in PD patients (for which bradykinesia is a hallmark symptom). We propose that the PMBR (and perhaps the MRBD) represent a continuum of movement speed increasing from HC eukinesia to HC slowed movement all the way to PD bradykinesia. Alternatively, this pattern may be best explained by the degree of motor plan complexity. Paradigms to intentionally slow movement can help distinguish the specificity of motor system disease biomarkers—to provide researchers/clinicians a tool to potentially differentiate physiological and pathophysiological beta activity.

## Data availability

The datasets generated during and/or analyzed from the current study are available by reasonable request from the authors.

